# An integrative multi-approach workflow resolves species limits in the southernmost members of the *Liolaemus kingii* group (Squamata: Liolaemini)

**DOI:** 10.1101/2020.07.02.185025

**Authors:** Kevin I. Sánchez, Luciano J. Avila, Jack W. Sites, Mariana Morando

## Abstract

Recent conceptual and methodological advances have enabled an increasing number of studies to address the problem of species delimitation in a comprehensive manner. This is of particular interest in cases of species whose divergence times are recent, where the conclusions obtained from a single source of evidence can lead to the incorrect delimitation of entities or assignment of individuals to species. The southernmost species of the *Liolaemus kingii* group (namely *L. baguali, L. escarchadosi, L. sarmientoi, L. tari* and the candidate species *L*. sp. A) show widely overlapping distributions as well as recent mitochondrial divergences, thus phylogenetic relationships and species boundaries are ambiguous. Here we use a comprehensive approach to assess species limits and corroborate their status as independent lineages through the use of four sources of molecular and morphological information (mitochondrial cytochrome-*b*, nuclear sequences collected by ddRADseq, and linear, meristic and landmark-based morphometrics). We found concordance among the different datasets, but signs of admixture were detected between some of the species. Our results indicate that the *L. kingii* group can serve as a model system in studies of diversification accompanied by hybridization in nature. We emphasize the importance of using multiple lines of evidence in order to solve evolutionary stories, and minimizing potential erroneous results that may arise when relying on a single source of information.

## 1. Introduction

Delimiting species objectively is critical in a wide range of biological research including the fields of evolution, biodiversity assessment and conservation. This may be a challenging task in recently derived populations due to the influence of the limited time since their split, which may heavily impact patterns of genetic differentiation (e.g. Nieto-Montes de Oca et al., 2017; Villamil et al., 2019). This is due to insufficient time for fixed differences to accumulate, which may compromise a consistent diagnosis (Struck et al., 2018), while admixture between closely related species can result in individuals with phenotypically and genetically intermediate states, generating controversial inferences with different data types (e.g. Sistrom et al., 2013; Olave et al., 2017; Singhal et al., 2018). On the other hand, there are documented cases of species with ancient divergence times that do not differ sufficiently in their morphological attributes, for example by niche conservatism (Wiens and Graham, 2005; Jezkova and Wiens, 2018), making difficult the assignment of individuals to putative species based solely on this class of characters. Processes such as past hybridization and incomplete lineage sorting of ancestral polymorphisms (ILS) can also mask the signatures of true phylogenetic divergence, and hence complicate the precise delimitation of species boundaries, especially, when relying on a single source of evidence (Maddison, 1997; Knowles and Carstens, 2007; Degnan and Rosenberg, 2009). Both of these processes have a high prevalence in recently diverged populations/species (Knowles and Carstens, 2007). But more recently, species delimitation research (SDL) has been characterised by a pluralistic approach in which independent sources of information (e.g. genetics, morphology, ecology) are used in order to objectively test species boundaries (Dayrat, 2005; Padial et al., 2010; Edwards and Knowles, 2014). Moreover, the increasing availability of genomic datasets comprised of hundreds/thousands of independent loci, and coalescent-based approaches for species delimitation, have opened the door for the detection of independent lineages with increased levels of statistical rigor and objectivity (e.g. Eaton and Ree, 2013; Leaché et al., 2014; Leaché and Oaks, 2017; Esquerré et al., 2019; Villamil et al., 2019; Dufresnes et al., 2020).

The lizard genus *Liolaemus* contains 271 species (Uetz, 2020) distributed in two main clades or subgenera: *Liolaemus sensu stricto* (the chilean group) and *Eulaemus* (the argentinean group; Laurent, 1983, 1985). The members of this genus are distributed throughout the Andes and adjacent lowlands of South America, from Perú to Tierra del Fuego in Argentina (Cei, 1986), having one of the largest latitudinal (14° ± 30’ - 52° ± 30’ S), elevational (from sea level to at least 5176 m.a.s.l.), and climatic distributions among lizards worldwide (Cei, 1986). This may explain the great diversity of biological characteristics observed in these reptiles, including body size (Pincheira-Donoso et al., 2015), karyotype (Morando, 2004; Aiassa et al., 2005), reproductive modes (viviparity/oviparity/parthenogenetic; Abdala et al., 2016), and dietary habits (insectivores/herbivores/omnivores; Cei, 1986, 1993; Espinoza et al., 2004). For this reason, this genus is increasingly being employed as a study model in numerous research areas such as evolution of herbivory, parity modes, community structure, comparative anatomy, physiology, behavioral ecology, and conservation (e.g. Espinoza et al., 2004; Tulli et al., 2009; Ibargüengoytía et al., 2010; Kacoliris et al., 2010; Corbalán et al., 2011; Kacoliris et al., 2011; Labra, 2011; Tulli et al., 2011). However, due to the species-richness and geographic range of the genus, taxonomy and systematics remain the most intensely studied focus (see Avila et al., in press, for a recent review).

Horizontal gene flow has been documented between non-sister species (e.g. *L. gracilis × L. bibronii*) (Olave et al., 2011, 2018), and many other instances of hybridization have been hypothesized with various levels of evidence for several groups within this genus. For example, Avila et al. (2006) have suggested the presence of hybridization between members of two deeply diverged species complexes (*L. boulengeri* and *L. rothi* complexes) based on extensive mitochondrial paraphyly, and more recently supported by morphological and nuclear sequence data analysed by recently developed coalescent-based methods to detect interspecific gene flow (Olave et al., 2018). For a detailed review of hybridization cases in *Liolaemus* see Morando et al. (2020).

The Patagonian lizards of the *Liolaemus kingii* group are included in the *L. lineomaculatus* section of the subgenus *Eulaemus* (Cei and Scolaro, 1982; Breitman et al., 2011, 2013), and currently includes 13 described species distributed in two main clades: the *L. kingii* clade (11 species) and the *L. somuncurae* clade (2 species) (Breitman et al., 2011, 2013; Olave et al., 2014). In addition, numerous candidate species have been identified based on various approaches, revealing that this group is characterized by high levels of genetic and morphological variation within and between populations, and phenotypic intergradation between adjacent populations (Breitman et al., 2013, 2015). The geographical distribution of this complex ranges mainly from southern Río Negro to southern Santa Cruz provinces in Argentina (41° ± 30’ - 52° ± 30’ S). Within the *L. kingii* clade, a widely distributed group of southern lineages, including *L. baguali, L. escarchadosi, L sarmientoi, L. tari* and the candidate species *L*. sp A identified by Breitman et al. (2015), is distributed across a heterogeneous area, ranging from sea level to high-altitude areas in the Andean cordillera, and characterised by recent divergence times (Late Pliocene to Late Pleistocene: ≈ 3 Mya to 0.75 Mya) (Breitman et al., 2011, 2013, 2015; Esquerré et al., 2018). These species were resolved as a well supported clade in a previous study attempting to infer relationships within the *L. lineomaculatus* section (Breitman et al., 2011), while other authors recovered conflicting topologies (see below). One challenging issue within this clade is the lack of objectivity in the species assignment for many individuals not collected from type localities. Further, this problem also extends to the *L. kingii* group as a whole, so the application of integrative taxonomic approaches will be helpful in future studies.

*Liolaemus sarmientoi* was described by Donoso-Barros (1973) from material previously adjudicated to a population of *L. dorbignyi* Koslowsky 1898, and included it within the “*L. kingii* group”. The type locality of this species was assigned to Monte Aymond, southern Santa Cruz province, Argentina (Donoso-Barros, 1973). Cei (1975, 1979) considered *L. sarmientoi* as a subspecies of *L. archeforus* Donoso-Barros and Cei 1971 (another member of the *L. kingii* group), and renamed the *“L. kingii* group” of Donoso-Barros (1973) as the “*L. kingii-archeforus* complex” (together with *L. archeforus archeforus* and *L. kingii* Bell 1843). In 1983, Cei and Scolaro described *L. baguali* (originally as *L. kingii baguali*), noting its distinctive color patterns and allopatric distribution with respect to *L. kingii*, and established Sierra del Bagual (southwestern Santa Cruz province, Argentina) as its type locality (Cei and Scolaro, 1983). Further, the taxonomic separation of the “*archeforus*” and “*kingii*” complexes (as defined in Cei and Scolaro, 1982) was reinforced and their geographical distributions restricted (Cei and Scolaro, 1983). Years later, Cei and Scolaro (1996) proposed that *L. baguali* and *L. sarmientoi* (and the other members of the *L. archeforus* and *L. kingii* groups) meet the criteria to be considered full species. Scolaro and Cei (1997) published a systematic review of the “*L. kingii*” and “*L. archeforus*” groups based on morphological attributes; they described *L. tari* and *L escarchadosi* (both included in the “*L. archeforus* group”), and highlighted their morphological resemblance to *L. sarmientoi*, with type localities in Del Viento Plateau and Cordón de los Escarchados, respectively (both in southwestern Santa Cruz province, Argentina). Breitman et al. (2011) could not verify the monophyly of the *L. kingii* and *L. archeforus* groups, and lumped all of these species into the *“L. kingii* + *archeforus* group” (Breitman et al., 2011). This clade was later renamed the “*L. kingii* group” by Breitman et al. (2013).

More recently, these species have been included in studies that aimed to evaluate phylogenetic relationships of members of the genus *Liolaemus*, or species groups within it, but conflicting topologies were inferred, mainly with the positions of *L. baguali* and *L. sarmientoi* (Espinoza et al., 2004; Schulte and Moreno-Roark, 2010; Breitman et al., 2011; Pyron et al., 2013; Olave et al., 2014; Breitman et al., 2015; Esquerré et al., 2018). In this context, the Breitman et al. (2015) study published was important because it focused specifically on these species and detected morphological evidence of hybridization between them. This may explain the ambiguity in the phylogenetic relationships between these species, but the boundaries between them have not been explicity tested with appropriate methods, so an empirical study is necessary to evaluate the results of different methods and datasets.

Herein, we used mitochondrial, genome-scale and morphological information to elucidate the phylogenetic relationships of these species, address species delimitation hypotheses and evaluate the extent of hybridization (Fig. 1). The use of different methodologies for phylogenetic inference and species delimitation, as well as different types of datasets, allowed us to assess both consistency across analytical methods and level of resolution provided by each source of information. Based on the problems mentioned above, we expected contrasting results with the different datasets employed and the different methodological approaches, which in turn are affected differentially by both shortcomings inherent to each method, and ongoing evolutionary processes.

**Figure 1.**
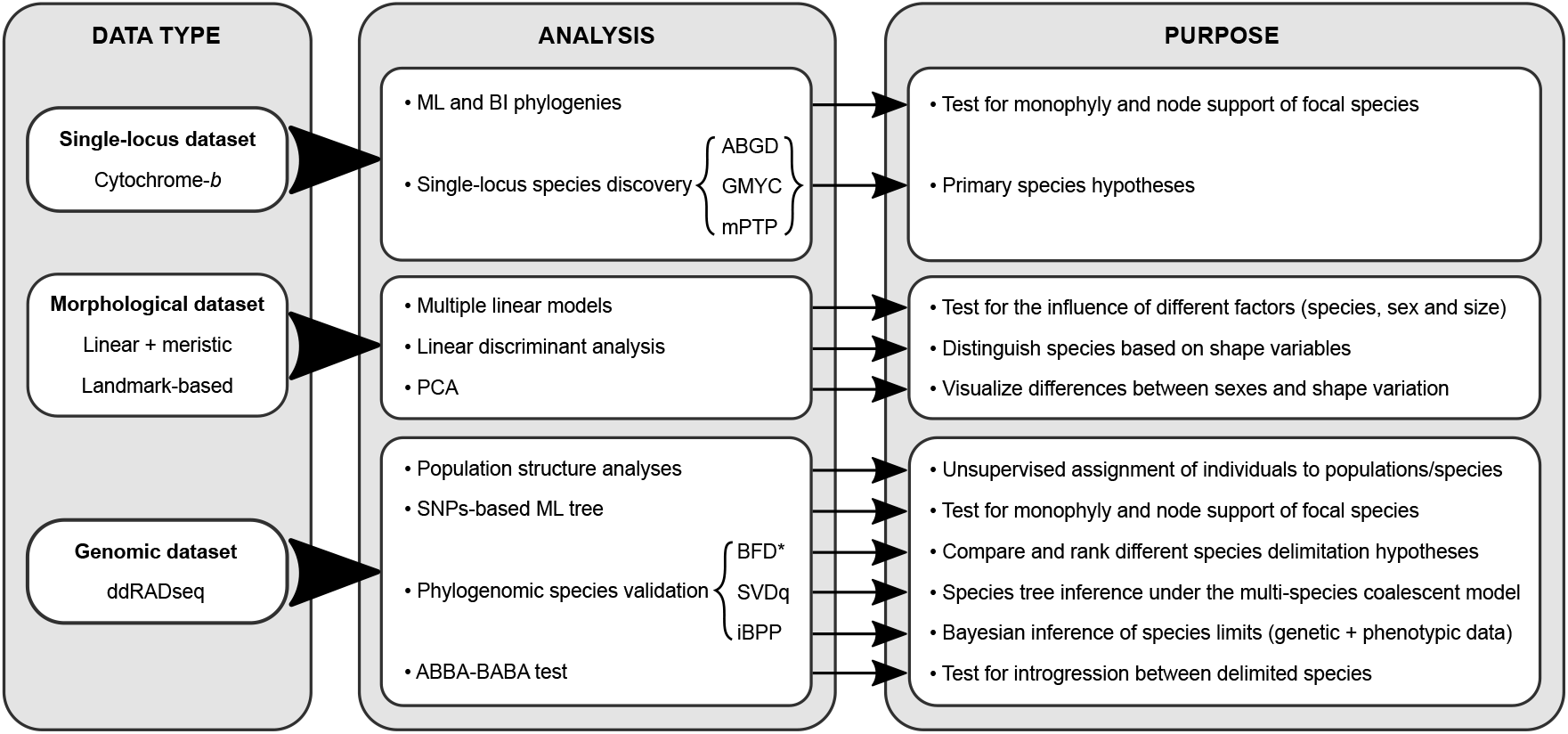
Summary of the integrative taxonomy workflow followed in this paper.

## 2. Materials and methods

### 2.1. Taxon sampling

We included samples of *L. baguali, L. escarchadosi, L. sarmientoi, L. tari* and the candidate species *L*. sp. A identified by Breitman et al. (2015). The nominal species were collected from type localities and across their known distributional ranges with the exception of *L. tari*, for which the closest sampling locality to the type locality was separated by ~ 25 km in SW direction (Fig. 2). Individuals were identified on the basis of their external morphological attributes and sampling location, according to their original descriptions. Sex was corroborated by the thickness of the base of the tail and the presence of precloacal pores (present mainly in males), and adults were identified by size and coloration patterns (Cei, 1986; Breitman et al., 2013). We included one sample of each of the remaining members of the *L. kingii* group for the mitochondrial tree, namely: *L. archeforus*, *L. chacabucoense*, *L. gallardoi*, *L. kingii*, *L. tristis*, *L. scolaroi*, *L. somuncurae*, *L. uptoni* and *L. zullyae*. For the ddRADseq-based tree, only *L. somuncurae* was included as outgroup. The voucher and locality data for each sample are provided in Appendix S1. Distribution maps were generated with QGIS v.3.10.1 (QGIS Development Team, 2020).

**Figure 2.**
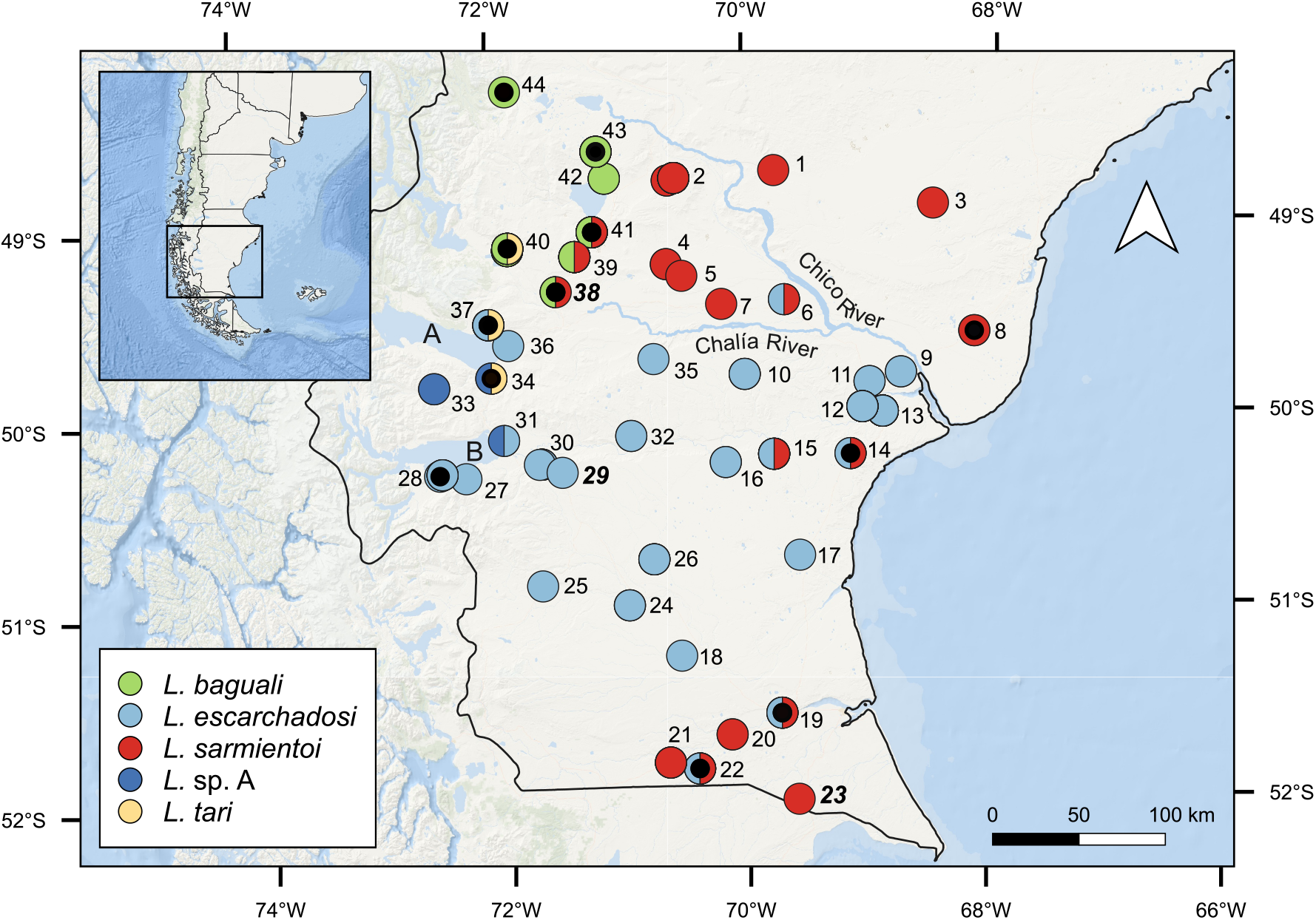
Sampled localities of the southern species of the *L. kingii* group. All points on the map are labeled with corresponding locality numbers, and type localities are indicated with bold-italic typefaces (23: type locality of *L. sarmientoi*; 29: TL of *L. escarchadosi* and 38: TL of *L. baguali*). Symbols are colored according to haplogroup memberships of the individuals based on the cyt-*b* gene tree. Smallest black circles represent localities for which individuals have ddRADseq data. A and B letters indicate the Viedma and Argentino lakes, respectively. Southern South America is depicted in the inset.

### 2.2. Data adquisition

#### 2.2.1. Mitochondrial data

We compiled 230 available sequences (224 ingroup plus 6 outgroup sequences) from GenBank, for the mitochondrial cytochrome-*b* gene from previous published studies on the *L. kingii* group (Breitman et al., 2011, 2015). Additionally, we generated new cyt-*b* data for 20 individuals (17 ingroup plus 3 outgroup sequences; GenBank accessions for each individual are provided in Appendix S1).

Genomic DNA was extracted from tissue (tail tips and liver) with either a Qiagen DNeasy blood and tissue extraction kit (Qiagen Inc., CA, USA) or NaCl-isopropanol extraction method (MacManes, 2013). The cytochrome-*b* region was amplified following the polymerase chain reaction and sequencing protocols of Morando et al. (2003).

Sequences were assembled and edited with Sequencher^®^ v.4.8 (Gene Codes Corporation, 2007) and aligned with the Muscle algorithm v.3.8.425 (Edgar, 2004), implemented in AliView v.1.26 (Larsson, 2014). Alignments were checked by eye to ensure there were no indels or stop codons and, if necessary, manually adjusted to maximize blocks of sequence identity. The total alignment consisted of 250 sequences of 813 bp.

Prior to species delimitation analyses (see below), we used RAxML v.8.2.10 (Stamatakis, 2014) to remove duplicate haplotypes from a matrix of 241 ingroup sequences. It is generally recommended that identical sequences be removed prior to tree-based methods of species delimitation (Blair and Bryson, 2017). This left a total of 115 haplotypes for species delimitation. Excluded specimens were subsequently assigned to putative species whose individuals shared identical haplotypes in order to perform morphological analyses (see below).

#### 2.2.2. Morphological data

Three types of morphological data were obtained: linear measurements (only in adults: 185 individuals), scale counts (juveniles and adults: 230 individuals), and dorsal head shape captured through geometric morphometrics (juveniles and adults: 226 individuals). We obtained 19 different linear measures to the nearest 0.1 mm using a Schwyz^®^ electronic digital calliper, plus 11 scale counts with a stereoscopic microscope (precloacal pores were recorded only on adults). See details on the definitions of linear measurements and scale counts in Appendix S2. All the measurements and counts were performed by the same person (KIS) to avoid error arising from inter-individual observations, and carried out on the right side of each specimen; we used the left side in the few cases data could not be collected from the right side (e.g. lack of a leg). Measurements and scale terminology followed Laurent (1986), Smith (1995), Etheridge (2000) and Breitman et al. (2013).

Geometric morphometric analyses were based on digital photographs of the dorsal view of the head of each specimen with a Lumix Fz 60 camera mounted in a fixed position. A rule was placed next to each one to scale the images. The characterization of the head shape was carried out placing 6 landmarks and 10 semilandmarks (Fig. S2) from the right side of each specimen, with the software tpsDig v.2.31 (Rohlf, 2015). For the open curve, all but the first and last landmarks were defined as sliding landmarks. As above, the digitizing step was performed by the same operator.

#### 2.2.3. Genomic data

We collected genomic data from 25 samples (24 ingroup and one outgroup sequences) with double-digest Restriction Site Associated DNA sequencing (ddRADseq; Peterson et al., 2012). We could not obtain sequences of *L*. sp. A, so the results based on the ddRADseq data are presented only for *L. baguali*, *L. escarchadosi*, *L. sarmientoi* and *L. tari*.

Libraries were prepared as follows: 1 mg of DNA from each individual was double-digested using SbfI and MspI enzymes. Fragments were then purified with Agencourt AMPure Beads and ligated with specific Illumina adaptors containing unique barcodes in order to associate reads to samples. Equimolar amounts of each sample were first multiplexed (pooled) and then size selected using PippinPrep with 2 % agarose gel cassettes (fragments of 100-600 bp were selected). Libraries were enriched using proofreading Taq and Illumina’s indexed primers. Enriched libraries were again purified with Agencourt AMPure Beads and quantified with PicoGreen before ligation of barcoded illumina adaptors onto the fragments. An Agilent Bioanalyzer was used to ensure that libraries were at the appropriate concentration and size distribution for sequencing. The sequencing was conducted on an Illumina HiSeq 2000 platform at the sequencing service of the University of California (Berkeley, USA) to generate single-end reads of 50 bp in length.

Sequence reads were demultiplexed and quality filtered with the *process_radtags* module of the Stacks v.2.41 pipeline (Rochette et al., 2019), using the following parameters: *-t 91, -c, -s 10, -r* and *-w 0.15*. Each locus was reduced from 50 to 39 bp after removal the barcode and adapter sequences. The pipeline ipyrad v.0.7.30 (Eaton and Overcast, 2020) was employed for additional filtering, clustering within and between individuals to identify homologous loci (full sequences, including invariable sites and single nucleotide polymorphisms), and exporting in different formats. The sequence similarity clustering threshold parameter of ipyrad (*clust_threshold*) was optimized quantitatively, assessing a range of values as suggested in McCartney-Melstad et al. (2019) looking at six metrics that include: (a) fraction of inferred paralogous clusters, (b) maximum proportion of SNPs and (c) heterozygosity recovered, (d) fraction of variance explained by the main principal components of genetic variation (e), relationship between missing data and genetic divergence, and (f) mean bootstrap values in a maximum likelihood tree building framework (Fig. S1). We assessed a range from 0.80 to 0.96 and, based on the optimization protocol, we selected a clustering threshold of 0.9 (90 % similarity for two sequences to be clustered together). In addition, we performed two assemblies modifying the minimum number of samples that must have data at a given locus for it to be retained in the final data set (*min samples locus* parameter): one with a minimum of 50 % of samples with data (*min_samples_locus = 12*) that include ingroup and outgroup sequences for phylogenetic analysis (named as *full dataset*: 3479 loci and 6505 SNPs), and another with 100 % of samples with data at any locus (*min_samples_locus = 24*) containing only ingroup samples (named as *ingroup dataset*: 752 loci and 1196 SNPs). This last dataset was assembled because some analyses do not require the inclusion of outgroups. Indeed, removing distant species should increase the amount of data recovered and the accuracy of the analyses (Pante et al., 2015). Since some analyses are more sensitive to missing data (e.g. population structure analysis, BFD*, see below), we performed additional filtering on the full and ingroup-only datasets with VCFtools v.0.1.16 (Danecek et al., 2011), including only biallelic sites and allowing for no missing data (parameters: --*min-alleles 2*, --*max-alleles* 2, --*maf 0.05* and --*max-missing 1*); this resulted in assemblies of 629 SNPs and 453 SNPs, respectively. These new assemblies were identified as the *full_filtered_dateset* and *ingroup filtered dateset*, respectively.

### 2.3. Single-locus genealogies and “de novo” species delimitation

We used maximum likelihood (ML) and Bayesian methods (BI) to infer gene trees from the cyt-*b* alignment. We performed the ML analysis using RAxML-HPC v.8 (Stamatakis, 2014) with the *GTRCAT* model of nucleotide substitution and the *-f a* option, which implements a rapid bootstrap analysis and searches for the best-scoring tree (Stamatakis, 2014). This analysis was run on the CyberInfrastructure for Phylogenetic RESearch (CIPRES Science Gateway v.3.3; Miller et al., 2010). A Bayesian time-calibrated tree was estimated with BEAST v.2.6.1 (Bouckaert et al., 2019) in conjunction with automatic substitution model selection implemented in bModelTest (Bouckaert and Drummond, 2017). For this analysis we selected a relaxed clock log-normal prior modeling a uniform distribution for the *clock.rate* parameter (mean = 0.0223, lower limit = 0.0143, upper limit = 0.0314, according to Fontanella et al., 2012), and a Yule tree prior for the speciation pattern. The calibration point was placed in the divergence between the *L. somuncurae* and *L. kingii* clades (both constituting the *L. kingii* group) following the time calibrated phylogeny of Esquerré et al. (2018) (normal distribution with mean = 6 and *σ* = 0.255 Mya). The analysis consisted of two independent runs for 1 × 10^8^ generations sampled every 5000 generations. Proper mixing of chains and convergence was assessed with Tracer v.1.7.1 (Rambaut et al., 2018), confirming minimum effective sample sizes (ESS) ≥ 300 for every parameter including tree topology. The tree files resulting from each independent run were merged with LogCombiner v.2.6.1 (Bouckaert et al., 2019), and a maximum clade credibility (MCC) keeping the target heights was annotated with TreeAnnotator 2.6.0 (Bouckaert et al., 2019). We interpreted bootstrap support (BS) values ≥ 70 and Bayesian posterior probabilities (PP) ≥ 95 % as evidence of significant support for a clade (Wilcox et al., 2002; Aguilar et al., 2016).

The species-delimitation analyses were performed in two sequential stages (Carstens et al., 2013; Pavón-Vázquez et al., 2018). The “de novo” species delimitation stage estimates the number of lineages (i.e. putative species) supported by molecular sequences or gene trees without *a priori* assignment of individuals to species, to generate a preliminary hypothesis of species limits (Carstens et al., 2013). Subsequently, in the “validation” step, the putative species from the previous step are joined or kept separate, confirming the previous delimitation or generating alternative hypotheses (Carstens et al., 2013).

The discovery (*de novo*) step was performed using one distance-based and three tree-based species delimitation methods: Automatic Barcode Gap Discovery (ABGD; Puillandre et al., 2012), maximum likelihood and Bayesian implementations of the Generalized Mixed Yule Coalescent Model (mlGMYC and bGMYC, respectively; Pons et al., 2006; Reid and Carstens, 2012; Fujisawa and Barraclough, 2013), and the multi-rate Poisson Tree Process (mPTP; Kapli et al., 2017).

ABGD infers a model-based confidence limit for intraspecific divergence based on prior intraspecific divergences, clustering similar haplotypes together as “species” (Puillandre et al., 2012). It requires that users provide several parameters: the choice of a distance metrics (Jukes–Cantor, Kimura-2P or p-distances), a prior limit to intraspecific diversity (P), and a proxy for the minimum gap width (X). For each set of user parameters, it returns two delimitations: a “primary partition” and a “recursive” one obtained after applying its algorithm recursively (Dellicour and Flot, 2018). Default settings were used for P (0.001, 0.1) and X (1.5), and testing the Kimura-2P, Jukes-Cantor and p-distances. The script *ABGDconsensus* (available at https://github.com/jflot/ABGDconsensus) was used to automate this approach and summarize the results in a single delimitation hypothesis.

The mlGMYC method works under a likelihood framework and delineates species by optimizing a mixed model that combines diversification between species (Yule model) and genealogical branching within species (coalescent model) and, in doing so, identifies the limit between the two processes as an abrupt increase in lineage accumulation at this point (Pons et al., 2006; Fujisawa and Barraclough, 2013). The Bayesian implementation (bGMYC) has the advantage of accounting for uncertainty in phylogenetic estimation (Reid and Carstens, 2012; Fujisawa and Barraclough, 2013). The R package Splits (SPecies’ LImits by Threshold Statistics;Ezard et al., 2009) was used to run the single-threshold mlGMYC, using the MCC tree obtained with BEAST 2. We assume a single threshold in mlGMYC due to better performance for simulated data compared to the multiple threshold approach (Fujisawa and Barraclough, 2013; Talavera et al., 2013; Luo et al., 2018). Next, we resampled 1000 random trees from the posterior distribution of BEAST 2 analysis and subjected them to the R package bGMYC (Reid and Carstens, 2012) to estimate the uncertainties in delimitation due to the phylogenetic estimation. The run consisted of 110,000 generations, discarding the first 10,000 as burn-in, and sampling every 100 generations afterwards. Because bGMYC returns a probability of conspecificity for each pair of sequences in the data set, its output is not directly comparable to the discrete grouping of individuals into putative species returned by other approaches (Dellicour and Flot, 2018). Here, we added a discretization step during which we retained only pairs of individuals that had a probability of conspecificity higher than 0.5, with the *bgmyc.point* function. We applied a threshold of 0.9, but only show the results for PP ≥ 0.5.

The mPTP method (Kapli et al., 2017) models intra and inter-specific processes (each as an independent Poisson process) based on the number of molecular substitutions on branches, so it does not require an ultrametric tree as input, instead, it requires that tree branch lengths be proportional to the number of substitutions. It has been shown that this method outperforms the original PTP model (Kapli et al., 2017) because each species branch has its own rate of evolution (*λ*) instead of sharing it, and is consistent and very effective for species delimitation in datasets with uneven sampling (Blair and Bryson, 2017). Using the best-scored ML tree from RAxML, we performed two simultaneous Markov Chain Monte Carlo (MCMC) runs of 1 × 10^8^ steps each, with a burn-in of 2 × 10^7^ and sampling every 10,000 steps. Convergence of the runs was assessed visually using the outputted likelihood plot of the combined runs (created using the --*mcmc_log* command). Outgroups were pruned before conducting the PTP analyses to avoid bias that may arise if some of the outgroup taxa are too distantly related to ingroup taxa (Zhang et al., 2013).

### 2.4. Morphological analyses

The linear measurements were transformed to log-shape ratios, obtained as the log of the division between the geometric mean of all measurements and each raw measurement (Mosimann, 1970; Claude, 2013). This approach loses one degree of freedom due to scaling (Claude, 2013). The effects of different factors on the variability of the linear measurements were tested. First, the effects of sex, species, and their interaction were estimated on geometric size (geometric mean) using a multiple linear model, and ANOVA with type II sums of squares was used to test the significance of each factor. Second, we applied a multiple and multivariate linear model on the first five principal components of shape variation (log-shape ratios) in order to test the effect of sex, species, and size factors, as well as the interactions between them. We performed a multivariate analysis of variance using type II sums of squares on the different variances and covariances explained by the factors and covariables. The individual assignment to species followed the results of the cyt-*b* gene tree. In addition, we performed a linear discriminant analysis using species assignment as group factor on (i) the log-shape ratios and (ii) the raw datasets, to evaluate whether it is possible to easily distinguish species based on these variables. The percentage of correctly assigned specimens was computed using a jackknife procedure. All these analyses were carried out in the R environment (R Core Team, 2019).

The analysis of the dorsal head shape data consisted of: (1) minimizing the bending energy of the semilandmarks with the tpsRelW v.1.70 software (Rohlf, 2015), and (2) scaling, translating, rotating and optimally superimposing the configurations through the Generalized Procrustes Analysis (Claude, 2008; Dryden and Mardia, 2016), with the *gpagen* function implemented in the R package geomorph v.3.2.0 (Adams et al., 2020).

Differences between sexes were explored using PCA for both datasets (log-shape ratios + meristic data and dorsal head shape data), and if sexes formed distinct clusters, we performed all subsequent analyses for males and females separately; otherwise data from both sexes were pooled. PCA was also employed to visualize how shape variation was structured and to identify which variables contribute most to the morphometric variation (Claude, 2008). For the log-shape ratios + meristic data, we used the function *PCA* implemented in the R package FactoMineR v.2.0 (Lê et al., 2008), and for dorsal head shape data we used the function *plotTangentspace* from geomorph.

### 2.5. Population structure analyses

Genetic structure was explored using the model-based fastSTRUCTURE software (Raj et al., 2014), which uses a variational Bayesian framework under a model assuming Hardy-Weinberg equilibrium and linkage equilibrium within populations. For this analysis we used the *ingroup filtered_dataset*, which was converted to the STRUCTURE format in PGDSpider v.2.1.1.5 (Lischer and Excoffier, 2012), and edited in a spreadsheet. We tested 1-10 genetic clusters by specifying the *k* parameter and running 5 replicates for each value with default convergence criterion and priors. The script *chooseK.py* was used to determine the value of *k* that best explained the structure underlying the data, and final plots were constructed using the script *distruct.py*, both included in fastSTRUCTURE. Additionally, we employed the model-free Discriminant Analysis of Principal Components (DAPC) implemented in the R package Adegenet v.2.0 (Jombart, 2008). This method makes no assumptions about population models, instead it defines synthetic variables in which genetic variation is maximized between clusters of individuals (*k*) and minimized within clusters. For this we transformed the *ingroup_filtered_dataset* into a *genind* object with the *import2genind* function. The *find.clusters* function was employed to evaluate *k* values ranging from 1 to 10, and the best-supported number of clusters was identified with Bayesian Information Criterion (BIC). Best *k* values were then considered for a Discriminant Analysis of Principal Components. Finally, membership probabilities of each individual to the DAPC clusters were inspected through the *compoplot* function of Adegenet. Because both methods lumped *L. escarchadosi* and *L. tari* (see Results), we performed an additional analysis considering only these two species in order to assess their distinctiveness.

### 2.6. Phylogenomic inferences and species validation

A maximum likelihood SNPs-based tree was inferred with RAxML-HPC v.8 (Stamatakis, 2014) in the CIPRES platform (Miller et al., 2010) using a *GTR+G+ASC_LEWIS* model. The ASC option was used to correct for an ascertainment bias in the likelihood calculations, given that SNPs only contain variable sites (Villamil et al., 2019).

We employed Bayes Factors Delimitation (BFD*; Leaché et al., 2014) implemented in the SNAPP template of BEAST 2 (Bryant et al., 2012), which allows a comparison of models that contain different numbers of species and different assignments of samples to species simultaneously. We used the assembly with no missing data (*ingroup-filetered_dataset*) to avoid overestimating the number of species (Leaché et al., 2018), and excluded samples with ≥ 5 % admixture in the population structure analyses, because horizontal gene flow is not taken into account in this method (see Results). This assembly was then converted to a nexus binary format in PGDSpider v.2.1.1.5 (Lischer and Excoffier, 2012). The evaluated models varied between two and five species and were based on: (A) cyt-*b* main haplogroups; (B) same as model A except that individual LJAMM-CNP 11582 was assigned to its own species following the results of GMYC and PTP; (C) same as model A except that *L. escarchadosi* and *L. tari* were lumped together based on results of ABGD; (D) results of DAPC which lumped *L. baguali* + *L. sarmientoi* on one hand and *L. escarchadosi* + *L. tari* on the other; (E) results of fast-STRUCTURE which lumped *L. escarchadosi* + *L. tari*; and (F) main lineages recovered in the SNPs-based tree (similar to model E except that *L. escarchadosi* and *L. tari* were considered different entities) (see details of each model in Table 3). The priors for the expected divergence (*θ*) were set according to the mean divergence within populations, using a gamma distribution with *α* = 1 and *β* = 70, and the priors for the speciation rate (*λ*) according to the maximum observed sequence divergence between any two taxa, using a gamma distribution with *α* = 2 and *β* = 200. To estimate the marginal likelihood of each species delimitation model, we conducted two independent path sampling analysis (*α* = 0.3) using 40 steps with an MCMC length of 300,000 generations, a pre-burnin of 30,000 and 10 % of final burn-in. This MCMC sampling frequency was sufficient to ensure that the majority of ESS values were ≥ 200. We ranked the models by their average marginal likelihoods (MLE) and calculated the Bayes Factors as BF = 2 × (MLE_1_ - MLE_0_), where MLE0 corresponds to model A and MLE1 to each alternative model (Kass and Raftery, 1995). A positive BF value indicates support in favor of model 1 while a negative BF value indicates support in favor of model 0. The strength of support for a model was obtained through the BF model selection statistic (calculated as 2 × ln(BF)), and assessed in the following manner: 0 ≤ 2 × ln(BF) ≤ 2 reflects weak support for model 1; 2 ≤ 2 × ln(BF) ≤ 6 reflects positive support; 6 ≤ 2 × ln(BF) ≤ 10 reflects strong support; and 2 × ln(BF) ≥ 10 reflects decisive support (Kass and Raftery, 1995).

We inferred a species tree under the multi-species coalescent model using the SVDquartets algorithm (Chifman and Kubatko, 2014) implemented in PAUP v.4.0a (Swofford, 2003). This method calculates the single-value decomposition (SVD) score for each possible four-species (quartet) topology in the dataset, and by optimizing the overall tree topology to maximize these scores of quartets induced by the tree topology (Chifman and Kubatko, 2014). The SVD score itself is calculated on the basis of a matrix of the genotype probabilities in a given quartet, and on the dimensionality of this matrix (see Chifman and Kubatko, 2014, for details). The great advantage of species-tree inference with SVDquartets is the very short time required to complete the analysis on a standard desktop computer. The input consisted of the unlinked SNPs output of ipyrad (*.u.snps*) with all the sequences included (*full dataset*). We performed one analysis with all specimens as independent samples and one where we assigned them to taxon partitions based on the best scored model inferred with SNAPP (see Results). Branch support was estimated by performing 1000 bootstrap replicates.

Finally, we implemented a joint Bayesian inference based on genetic and phenotypic data to delimit species using iBPP v.2.1.3 (“integrative Bayesian Phylogenetics and Phylogeography”; Solís-Lemus et al., 2015). This method was built upon the same Bayesian framework of the BPP software (Yang, 2015) to model molecular data across multiple genes, and it assumes a known assignment of individuals to putative species and a guide tree of these putative species. It has the advantage of incorporating independent quantitative continuous traits, which are modeled by a Brownian motion (BM) process along the species tree (Solís-Lemus et al., 2015). Alternative hypotheses about delimited taxa are generated by collapsing one or more nodes in the guide tree, each of which is given an equal prior probability. The posterior probability of a particular number of taxa, as specified by a given species tree and its branch lengths, is calculated from independent gene trees according to the multispecies coalescent (MSC) (Rannala and Yang, 2003). We based our individual assignment to species on the best-scored model inferred with SNAPP, and the guide tree on the result of SVDquartets (see Results). Three different datasets were employed: (1) molecular and morphological data combined, (2) molecular data alone and (3) morphological data alone. This method assumes gene flow ceases immediately following speciation, so individuals who showed signs of admixture (≥ 5 %) were removed. The morphological character matrix included the first five principal components of variation for the dorsal head shape data plus the first four components of the meristic data, and the molecular matrix consisted of the *.loci* output file from ipyrad, transformed to a PHYLIP assembly with the ipyrad-analysis toolkit (Eaton and Overcast, 2020). Following Huang and Knowles (2016) we assessed support for putative taxa with different demographic scenarios. For this we performed four analyses, each specified with a distinct combination of gamma priors for theta (*θ*) and tau (*τ*) parameters modelling: (1) large ancestral population sizes and deep divergences (*θ* = *G*(1, 10), *τ* = *G*(1, 10)), (2) large ancestral population sizes and shallow divergences (*θ* = *G*(1, 10), *τ* = *G*(2, 2000)) (3) small ancestral population sizes and deep divergences (*θ* = *G*(2, 2000), *τ* = *G*(1, 10)) and (4) small ancestral population sizes and shallow divergences (*θ* = *G*(2, 2000), *τ* = *G*(2, 2000)). Each analysis consisted of two replicates. The priors for the trait data were set as default values (*σ*^2^ and *κ* = 0). For other divergence-time parameters we used the Dirichlet prior (Yang and Rannala, 2010). We used the standard species delimitation algorithm (*uniformrootedtrees = 1*) that assigns equal probabilities to rooted species trees, since the alternative algorithm (*uniformrootedtrees = 0*) assigns equal probabilities to all labeled histories and can over-resolve large unbalanced guide trees (Yang and Rannala, 2010). We choose an automatic adjustment for the MCMC steps length, and the analyses consisted of runs of 6 × 10^4^ generations and a burn-in of 1 × 10^4^. Posterior probabilities ≥ 0.95 were considered as support for a speciation event. The control files (*.ctl*) were generated with *BPPmulti* (perl script available at http://github.com/eberlejonas/BPPmulti).

### 2.7. Tests for introgression with genomic data

We calculated the Patterson’s *D*-statistic (or ABBA-BABA test) to test for introgression after the point of lineage divergence for all species pairs (Green et al., 2010; Durand et al., 2011; Eaton and Ree, 2013). This test uses biallelic SNPs based on a four-taxon pectinate tree with the form (((P1,P2),P3),O), where the Outgroup taxon is included to determine which allele is ancestral (the “A” allele) and which is derived (the “B” allele). The *D*-statistic is calculated based on the number of patterns that conflict with the tree (the “ABBA” and “BABA” patterns) as *D* = (nABBA - nBABA)/(nABBA + nBABA). Under incomplete lineage sorting alone, two sister species (P1 and P2) are expected to share about the same proportion of derived alleles with a third closely related species (P3). Then, the number of derived alleles shared by P1 and P3 but not P2 and the number of derived alleles shared by P2 and P3 but not P1 should be approximately similar, resulting in *D* ≈ 0 (i.e. (A, (B, (B, A))) ≈ (B, (A, (B, A)))). In contrast, if hybridization leads to introgression between species P3 and either P1 or P2, then P3 should share more derived alleles with that species than it does with the other, leading to asymmetry in the sharing of derived alleles and an excess of either “ABBA” or “BABA” site patterns. To the extent that RAD sequences are a random sample of the genome, the *D*-statistic represents a genome-wide measure of introgression (Eaton and Ree, 2013). The *Dtrios* routine implemented in the software DSuite v.0.3 (Malinsky, 2019) was employed to calculate *D*-statistics using the *full_filtered_dataset*. This software keeps the outgroup fixed and tests all possible ways in which three species can be selected from the ingroup species and placed into the positions P1, P2, and P3; significance was evaluated with a *p*-value based on jackknifing against the null hypothesis of *D* = 0 (Malinsky, 2019). *P*-values were adjusted to account for the multiple comparisons issue (Malinsky, 2019) with the Benjamini-Yekutieli correction (Benjamini and Yekutieli, 2001), implemented in the function *p.adjust* in R. The individual assignments to putative species followed the results of fastSTRUCTURE, and the SVDquartets tree was used to specify species relationships. In addition, we reconstructed an unrooted phylogenetic network with the NeighborNet algorithm in SplitsTree4 (Huson and Bryant, 2006) using the *ingroup_filetered_dataset* as input. Bootstrap analyses were performed using 1000 replicates to generate the consensus network.

## 3. Results

### 3.1. Single-locus genealogies and “de novo” species delimitation

With cytochrome-*b*, the MCC and ML trees inferred five main lineages corresponding to *Liolaemus baguali*, *L. escarchadosi*, *L. sarmientoi*, *L. tari* and the candidate species *L*. sp. A (Fig. 3). *Liolaemus sarmientoi* represented the earliest-diverging lineage and was more closely related to the remaining members of the *L. kingii* clade than to the clade composed of *L. baguali*, *L*. sp A, *L. tari* and *L. escarchadosi*. This latter clade was inferred in a pectinate topology and the main haplogroups had high support with both reconstruction methods. An expanded version of the MCC tree with voucher labels is presented in Fig. S3.

**Figure 3.**
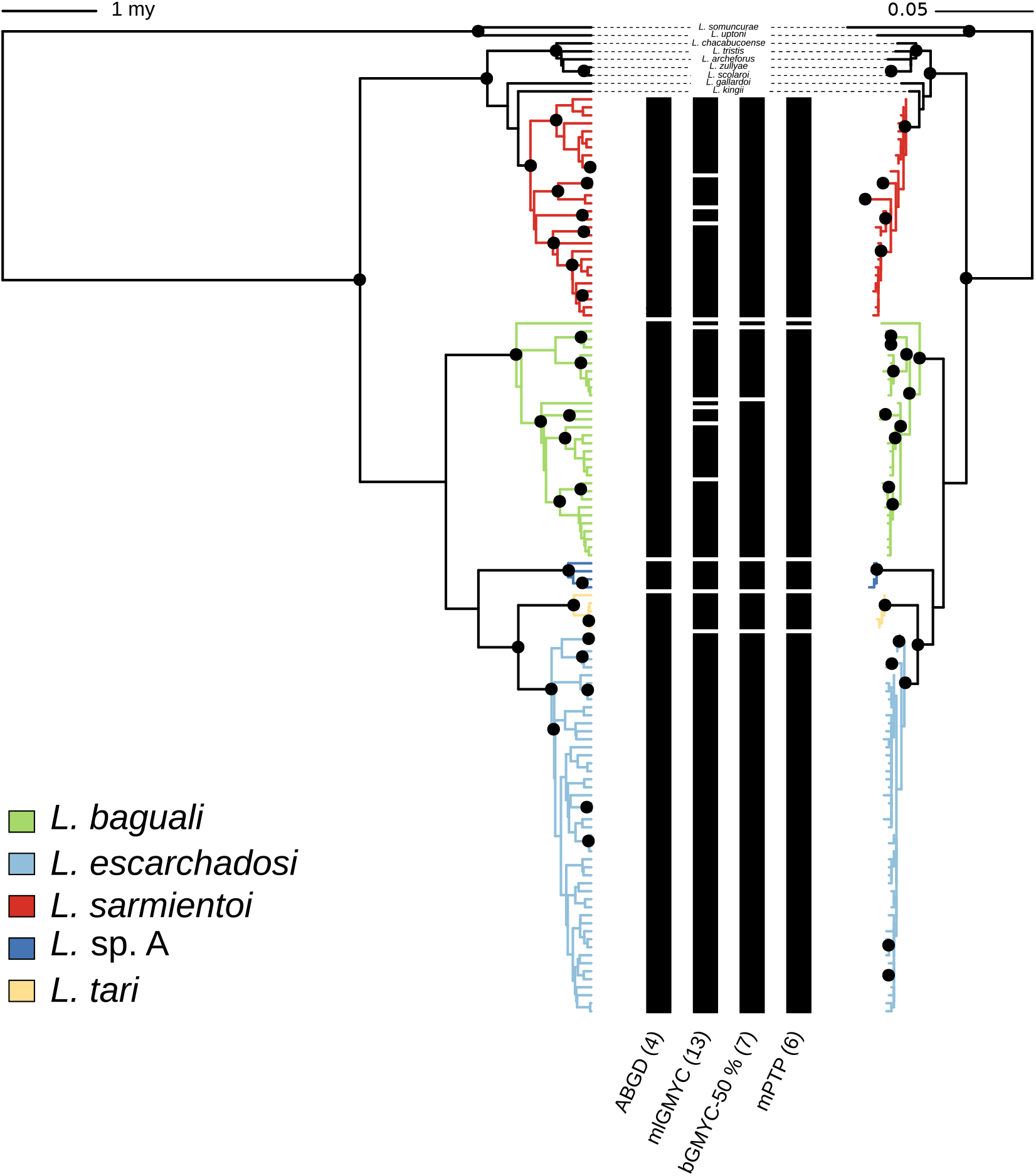
Cytochrome-*b* gene trees with summary of single-locus species delimitation results. The partition scheme obtained for each method is indicated by black boxes, and the number of delimited entities is detailed in parenthesis next to the name of each method. Gene trees correspond to ultrametric time-calibrated Maximum Clade Credibility tree inferred with BEAST 2 (left), and maximum likelihood tree inferred by RAxML (right). Black nodes indicate posterior probabilities (PP) ≥ 0.95 and bootstrap support ≥ 70, respectively.

Of the four *de novo* species delimitation approaches, ABGD was the most conservative since it found support for four of the five haploclades inferred in the gene trees, and lumped *L. escarchadosi* and *L. tari* into a single entity (Fig. 3). In contrast, the maximum-likelihood implementation of the GMYC model (mlGMYC) estimated 13 entities with a wide confidence interval of 4-34 (LRT = 0.00604108**), splitting *L. sarmientoi* into four entities (clades) and *L. baguali* into six entities (four clades plus two singletons). The Bayesian implementation (bGMYC) suggested seven entities when the probability of conspecificity threshold was set to 0.5, and these were composed of the *L. sarmientoi*, *L. tari*, *L*. sp A, and *L. escarchadosi* haplogroups plus *L. baguali*; the latter was split into two clades and one singleton (LJAMM-CNP 11582). Further, this singleton has the most northern distribution of this species, and it is also recognized in mlGMYC as an independent entity (see Fig. 2). When the probability threshold was set to 90 % (0.9) the number of independent entities scaled to 20, lumping predominantly specimens from the same sampling locality (results not shown). The MCMC run of the mPTP method retrieved similar results to ABGD except for the recognition of *L*. sp A and the haplotype LJAMM-CNP 11582 of *L. baguali* as independent lineages (see Fig. 3).

### 3.2. Morphometric analyses

Results of the linear models indicate that size was significantly related to sex dimorphism and species, but the interaction between these factors was not significant (Table 1). On the other hand, sex, species and size significantly explained shape variation (given by log-shape ratios); likewise, the interactions sex:size and species:size also proved to be significant (Table 2).

**Table 1:** ANOVA on geometric size (geometric mean). *P*-values below significance levels are shown in bold

| Factor | Sum of squares | df | *F*-test | *p*-value |
| --- | --- | --- | --- | --- |
| sex | 15.354 | 1 | 30.3888 | **$1.25 \times 10^{-7}$** |
| species | 15.354 | 4 | 4.1520 | **0.003079** |
| sex:species | 0.956 | 4 | 0.4729 | 0.755572 |
| Residuals | 88.419 | 175 |  |  |

**Table 2:**
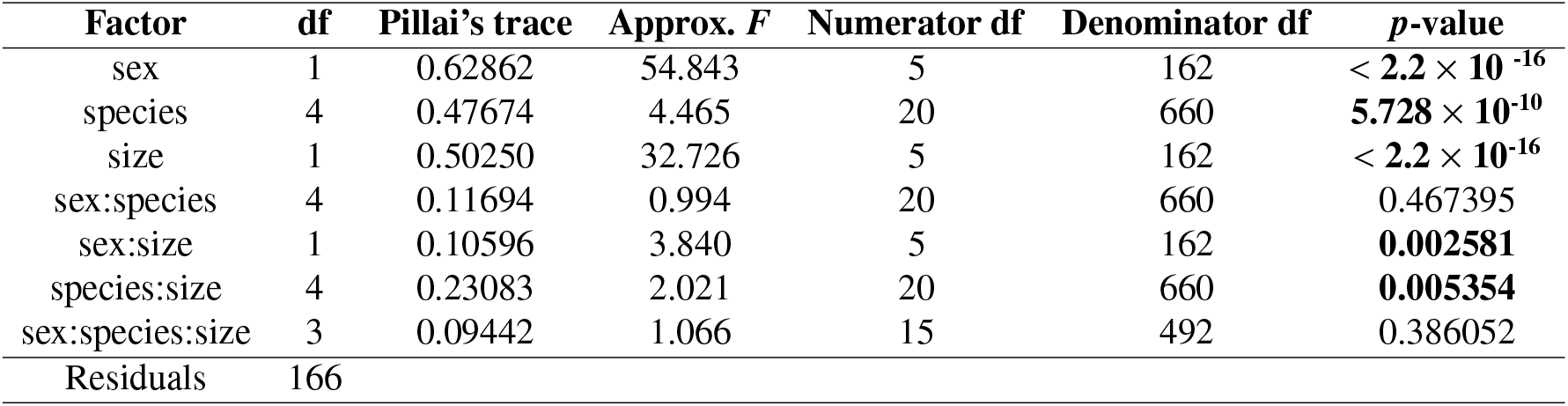
MANOVA on the first five PCs of log-shape ratios. *P*-values below significance levels are shown in bold

The predictive discriminant analysis based on specimens identified by the cyt-*b* haplogroup correctly assigned only 55 % (0.5513514) of the individuals, this low percentage probably due to unbalanced sampling (Claude, 2013). When a discriminant analysis was directly applied to body shape measurements (thereby including size), the percentage of correctly classified individuals reached 63 % (0.6324324); hence, by introducing this new variable, the linear combination of variables corresponding to discriminant coefficients was slightly more efficient to discriminate between species.

Overall, the PCA analysis showed large overlap in the morphological variation, mainly in dorsal head shape, obscuring most putative species boundaries (Figs. 4 and 5). The first three principal component axes on body shape (log-shape ratios + meristic data) explained only 17.24 %, 11.43 % and 8.01 % of the total variation, respectively (Fig. 4; Table S1). PC_1_ was mostly loaded by AL, FL, IS, SAM, DS, VS, IL3 and IL4 in *L. baguali* (positive), and RW in *L. escarchadosi* (negative). The remaining lineages showed high overlap with the exception of *L*. sp A. This lineage could be differentiated from the rest of the species in PC_3_, which was positively loaded by HL and negatively by AGD and BL (Fig. 4). On the other hand, the first three PCs on variability in dorsal head shape explained only 52.41 % (PC_1_ = 25.41 %, PC_2_ = 14.53 % and PC_3_ = 12.47 %), indicating that this attribute is highly conserved in these species (Fig. 5; Table S2).

**Figure 4.**
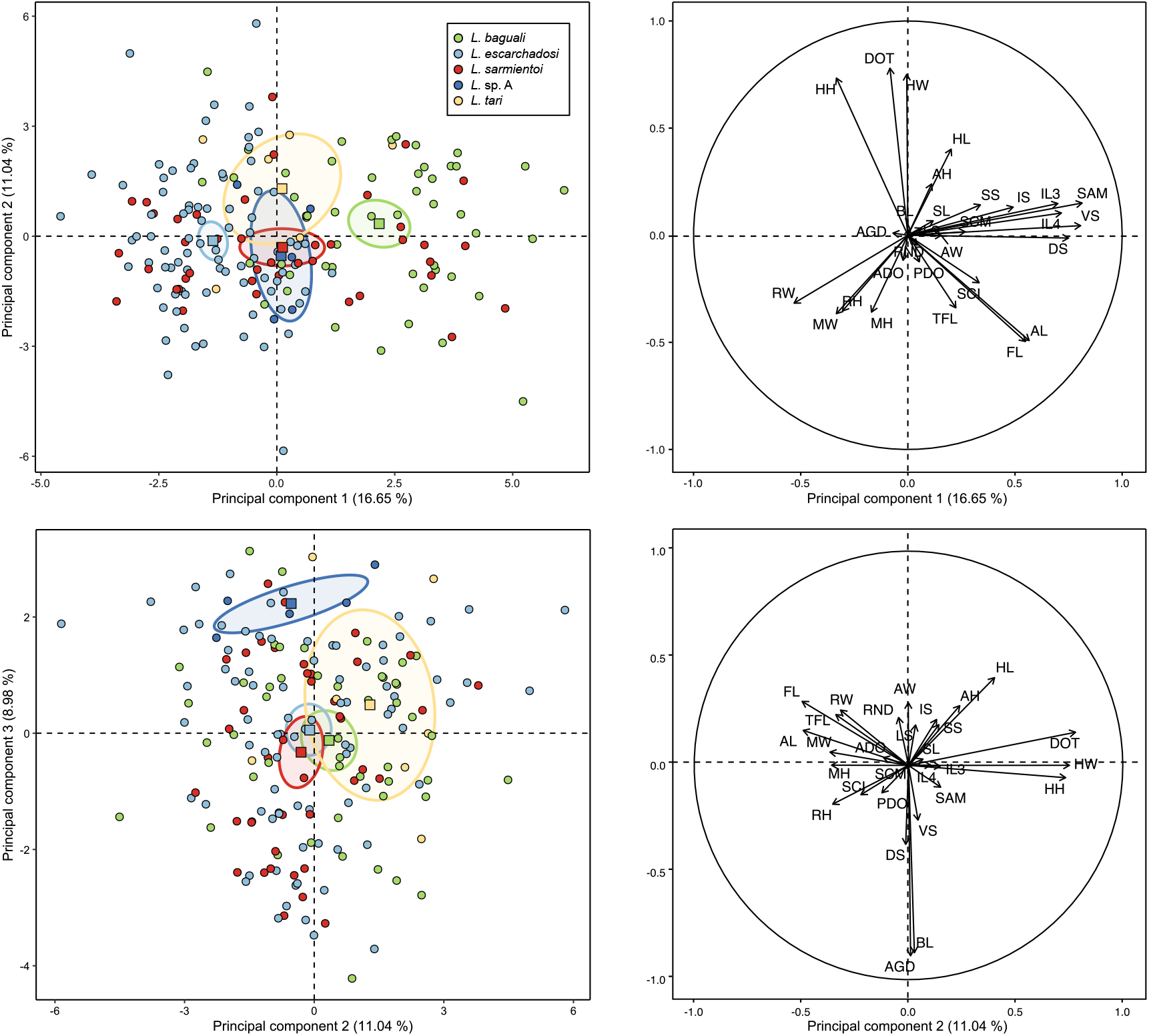
Principal component analysis plots of linear measurements (log-shape ratios) plus meristic variables. On the left plots of PC_1_ vs PC_2_ and PC_2_ vs PC_3_ with ellipses of 95 % confidence around each species barycenter. On the right variable factor maps depicting the variable loadings for each PC.

**Figure 5.**
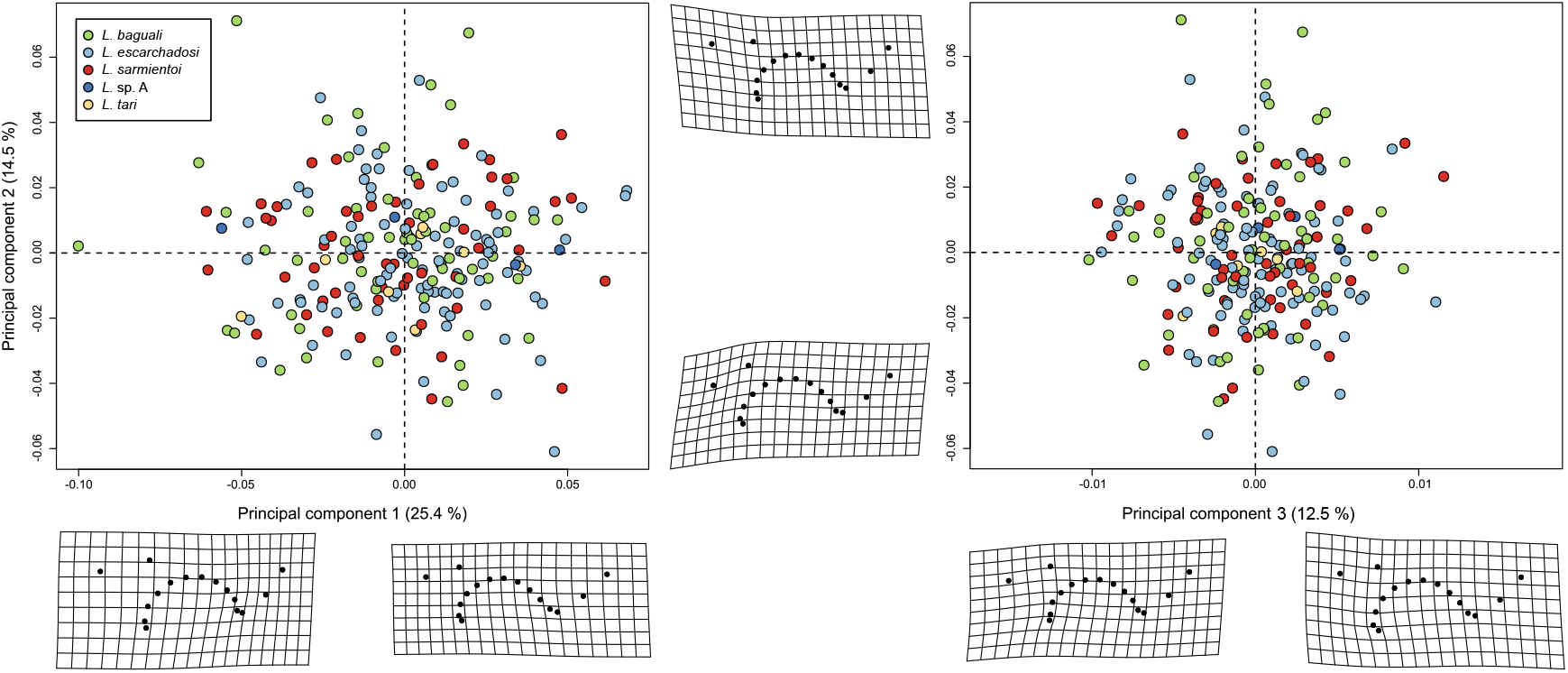
Principal component analysis plots of dorsal head shape variability and deformation grids illustrating the configurations at the extreme of each axis. PC_1_ vs PC_2_ plot on the left and PC_2_ vs PC_3_ plot on the right. Note the high overlap in the distribution of the points

### 3.3. Population structure analyses

The model-based fastSTRUCTURE indicated that a value of *k* = 2 maximized the marginal likelihood while a value of *k* = 3 best explained the structure observed in the dataset, thus, we present the results for *k* = 2 and *k* = 3. The two population clusters model (k = 2) corresponded to the lumping of *L. sarmientoi, L. baguali* and the specimen LJAMM-CNP 9349 assigned to *L. tari* in the cyt-*b* gene tree in one group, and the remaining members of *L. tari* plus *L. escarchadosi* in the other. The three clusters model (*k* = 3) corresponded to *L. sarmientoi, L. baguali* + LJAMM-CNP 9349, and the remaining members of *L. tari* + *L. escarchadosi*. In addition, one member of *L. sarmientoi* and two members of *L. tari* showed signs of admixture (LJAMM-CNP 11450, LJAMM-CNP 7253 and LJAMM-CNP 9349, respectively) (Fig. 6a, b). With DAPC, the *k*-means clustering algorithm revealed the lowest BIC value for two clusters, while values for more clusters were higher (Fig. S4). This result corresponded to those of fastSTRUCTURE for *k* = 2. The main difference was that DAPC clearly separated the groups in the genomic space with no signs of admixture, assigning each individual with 100 % probability to their respective “species” (Fig. 6d). *Liolaemus tari* and *L. escarchadosi* were clearly distinguished with both methods when they were analyzed separately, revealing a fine structure not detected when we included all the species (Fig. 6c, e).

**Figure 6.**
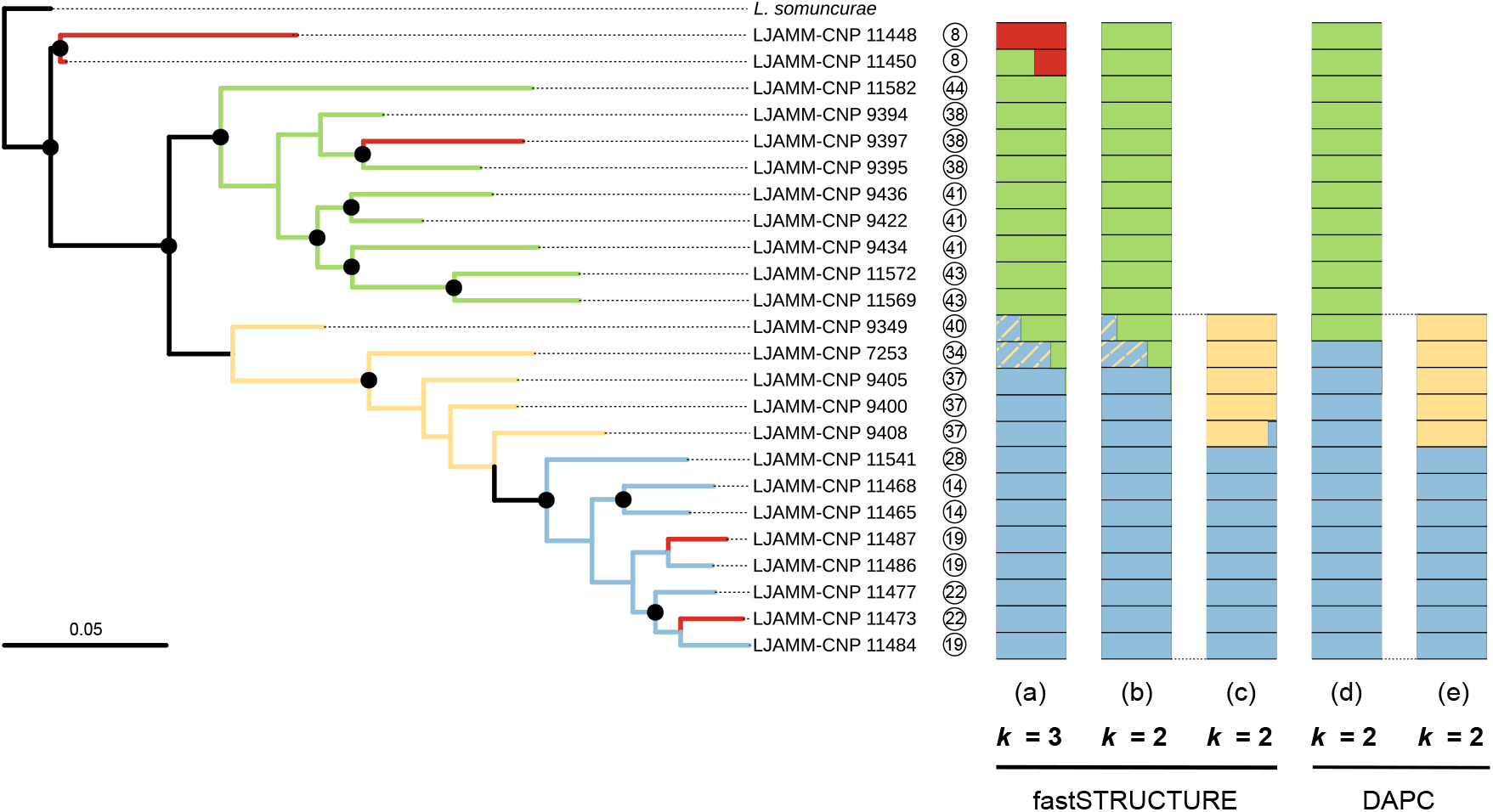
Left: maximum likelihood tree based on 6505 SNPs; terminals are colored according to the cyt-*b* tree. Right: corresponding plots of cluster membership inferred with fastSTRUCTURE and DAPC. Numbers within circles indicate the sampling locality for each individual. The stripped yellow lines within the distruct plots corresponding to specimens LJAMM-CNP 9349 and LJAMM-CNP 7253 highlight the members of *L. tari* that showed admixed genomic compositions

### 3.4. Phylogenomic inferences and species validation

The ddRADseq-based ML tree inferred the same relationships obtained with the cyt-*b* gene tree: a clade composed of two individuals of *L. sarmientoi* appeared as a basal lineage and sister to the clade composed of the remaining species, which in turn were inferred with high support except for *L. tari* (Fig. 6). However, some discordances were found between these two datasets. First, three specimens inferred as *L. sarmientoi* in the cyt-*b* gene tree were inferred with high support as the sister group to members of other species in the ddRADseq tree: one female specimen within the *L. baguali* clade (LJAMM-CNP 9397) and two specimens nested within the *L. escarchadosi* clade (male LJAMM-CNP 11473 and female LJAMM-CNP 11487). Second, *L. tari* was inferred as paraphyletic with respect to *L. escarchadosi* in the ddRADseq tree (see Fig. 6).

The results for the six models tested with BFD* are summarised in Table 3. Model F, which considers *L. escarchadosi* and *L. tari* as separate species and assigns individuals based on the ddRADseq tree, was the top-ranked model; it has the highest MLE value and was supported in favor of the null hypothesis that assigned individuals based on the cyt-*b* haplogroups (model A). The Bayes Factor in support for model F was decisive compared to model A.

**Table 3:** Stepping stone sampling results for the six species delimitation models. *L. ba.: Liolaemus baguali; L. es.: L. escarchadosi; L. sa.: L. sarmientoi; L. ta.: L. tari*; MLE: maximum likelihood estimation; BF: Bayes factor

| N species | Model | MLE | BF | $2 \times \ln(\text{BF})$ | Rank |
| --- | --- | --- | --- | --- | --- |
| 4 | A: (*L. sa.*) (*L. ba.*) (*L. ta.*) (*L. es.*)<sup>1</sup> | -8806.658 | - | - | 6 |
| 5 | B: (*L. sa.*) (LJAMM-CNP 11582) (*L. ba.*) (*L. ta.*) (*L. es.*)<sup>1</sup> | -8765.479 | 82.357 | 8.822 | 4 |
| 3 | C: (*L. sa.*) (*L. ba.*) (*L. ta.* + *L. es.*)<sup>1</sup> | -8903.588 | -193.861 | 10.534 | 5 |
| 2 | D: (*L. sa.* + *L. ba.*) (*L. ta.* + *L. es.*)<sup>2</sup> | -8473.267 | 666.782 | 13.005 | 3 |
| 3 | E: (*L. sa.*) (*L. ba.*) (*L. ta.* + *L. es.*)<sup>2</sup> | -8397.661 | 817.993 | 13.414 | 2 |
| 4 | F: (*L. sa.*) (*L. ba.*) (*L. ta.*) (*L. es.*)<sup>2</sup> | -8256.658 | 1099.999 | 14.006 | 1 |

<sup>1</sup>Specimens LJAMM-CNP 9397, LJAMM-CNP 11473 and LJAMM-CNP 11487 assigned to *L. sarmientoi*, according to the *cyt-b* gene tree
<sup>2</sup>Specimen LJAMM-CNP 9397 assigned to *L. baguali* and specimens LJAMM-CNP 11473 and LJAMM-CNP 11487 to *L. escarchadosi*, according to the ddRADseq-based tree

When all specimens were considered as independent samples, the SVDquartets tree reflected the same relationships evidenced in the ML ddRADseq tree (Fig. 7). However, the clade (*L. tari* + *L. escarchadosi*) was inferred with high support, and there were some differences in the internal relationships within the main clades. The taxon partition based on the top ranked model of SNAPP recovered the branches that separate *L. baguali* from (*L. escarchadosi* + *L. tari*), and *L. escarchadosi* from *L. tari* with high support (BS = 100) (Fig. 7).

**Figure 7.**
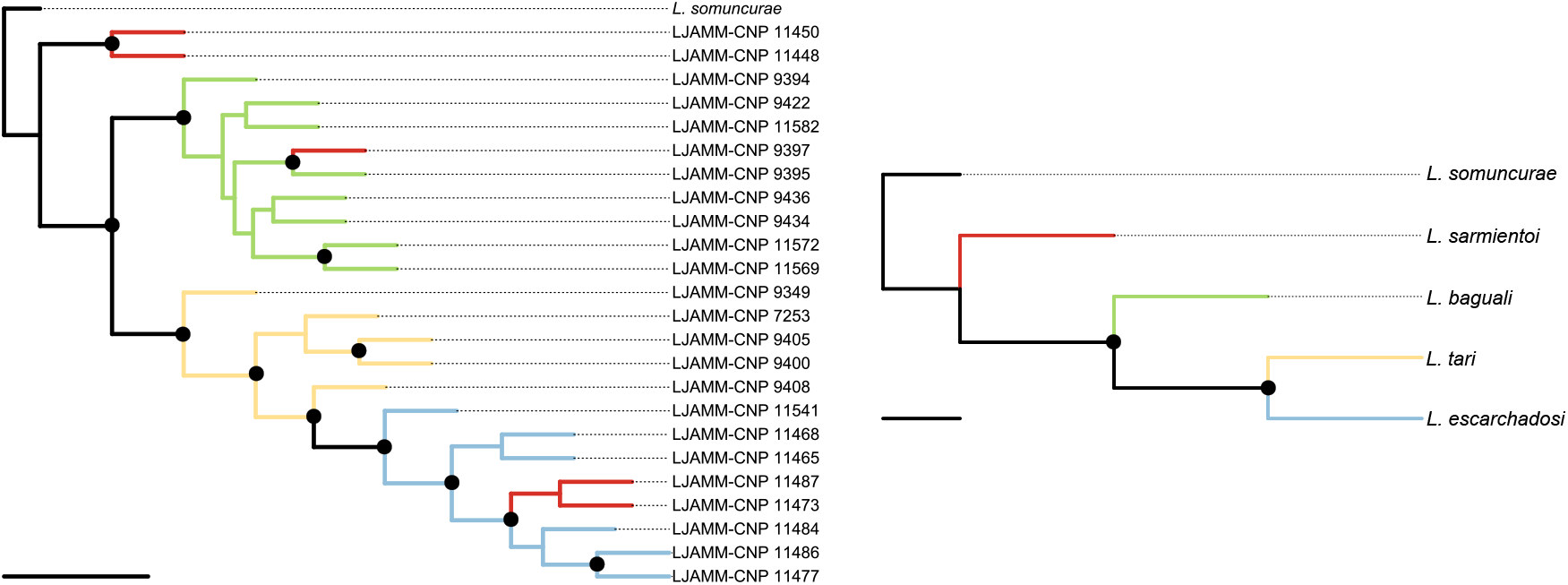
Species tree based on 2750 unlinked SNPs inferred with SVDquartets with all specimens as independent samples (left), and with taxon partitions based on population structure analyses (right). In the first tree the terminals are colored according to the cyt-*b* tree. Black nodes indicate Bootstrap Support ≥ 70.

Results from iBPP on the guide tree that considers *L. escarchadosi* and *L. tari* as independent lineages based on the best-scored model inferred with SNAPP were concordant with the four prior combinations. All the nodes were inferred with high support (PP ≥ 0.95) (Fig. 8). The analyses based on genetic and morphological data alone inferred similar posterior probabilities compared with the joint analysis of genetic and morphological data, irrespective of the differences in the demographic parameters tested (Fig. 8).

**Figure 8.**
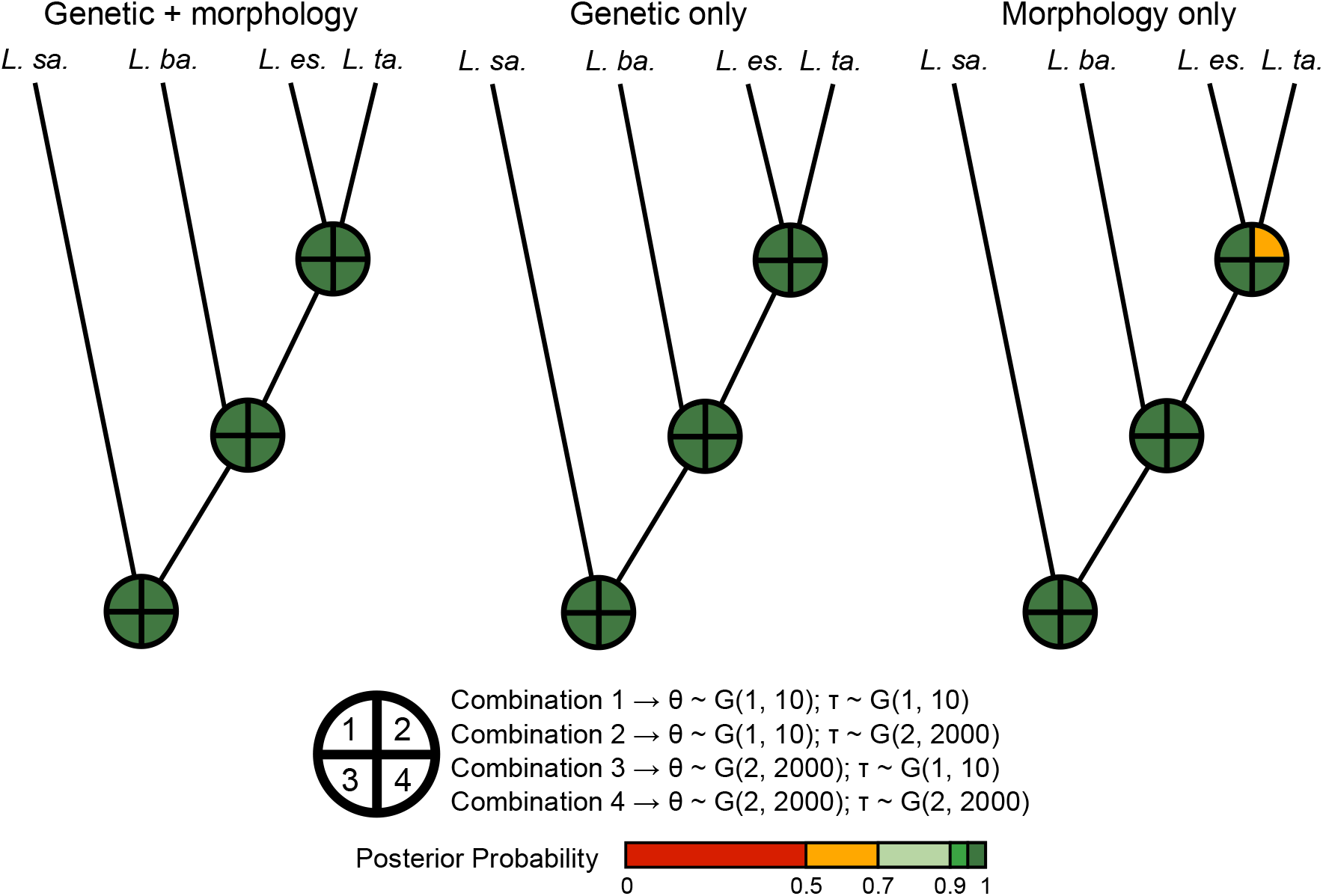
Results of species delimitation analyses with iBPP. The colored pie at each node represent the mean posterior probability for each gamma prior combination (*G*(*α,β*)) for ancestral population size (*θ*) and root age (*τ*). *L. sa: L. sarmientoi; L. ba.: L. baguali; L. es*.: *L. escarchadosi*; *L. ta*.: *L. tari*

### 3.5. Tests for introgression with genomic data

The *D*-statistics calculated for all the trios involving *L. baguali*, *L. escarchadosi*, *L. sarmientoi* and *L. tari* were greater than zero, indicating an excess of ABBA sites in all cases, although the values were small (0.05-0.17). All the tests showed significant values except for the trio *L. tari* (P1), *L. baguali* (P2) and *L. sarmientoi* (P3) (Table 4; Fig. S5). This pattern suggests weak nuclear introgression between *L. baguali* and *L. sarmientoi*, *L. tari* and *L. baguali*, and *L. tari* and *L. sarmientoi*. The SplitsTree network evidenced a wide reticulation in the base of the branches but some genetic clusters that match taxonomic treatments could be distinguished (i.e.: *L. sarmientoi*, *L. baguali* and *L. escarchadosi*) (Fig. 9). *Liolaemus tari* individuals appeared in the middle between these main groups without a clear distinction, which is in agreement with the population structure analyses (Fig. 6).

**Table 4:** Summary of ABBA-BABA tests. *Liolaemus somuncurae* was used as outgroup for all tests. *P*-values below significance levels are shown in bold

| **P1** | **P2** | **P3** | ***D*-statistic** | **Uncorrected *p*-value** | **Corrected *p*-value** |
| --- | --- | --- | --- | --- | --- |
| *L. escarchadosi* | *L. baguali* | *L. sarmientoi* | 0.164014 | 0.00601835 | **0.016717639** |
| *L. escarchadosi* | *L. tari* | *L. baguali* | 0.172074 | 0.00115408 | **0.009617333** |
| *L. tari* | *L. baguali* | *L. sarmientoi* | 0.0566751 | 0.204417 | 0.42586875 |
| *L. escarchadosi* | *L. tari* | *L. sarmientoi* | 0.178139 | 0.00263276 | **0.010969833** |

**Figure 9.**
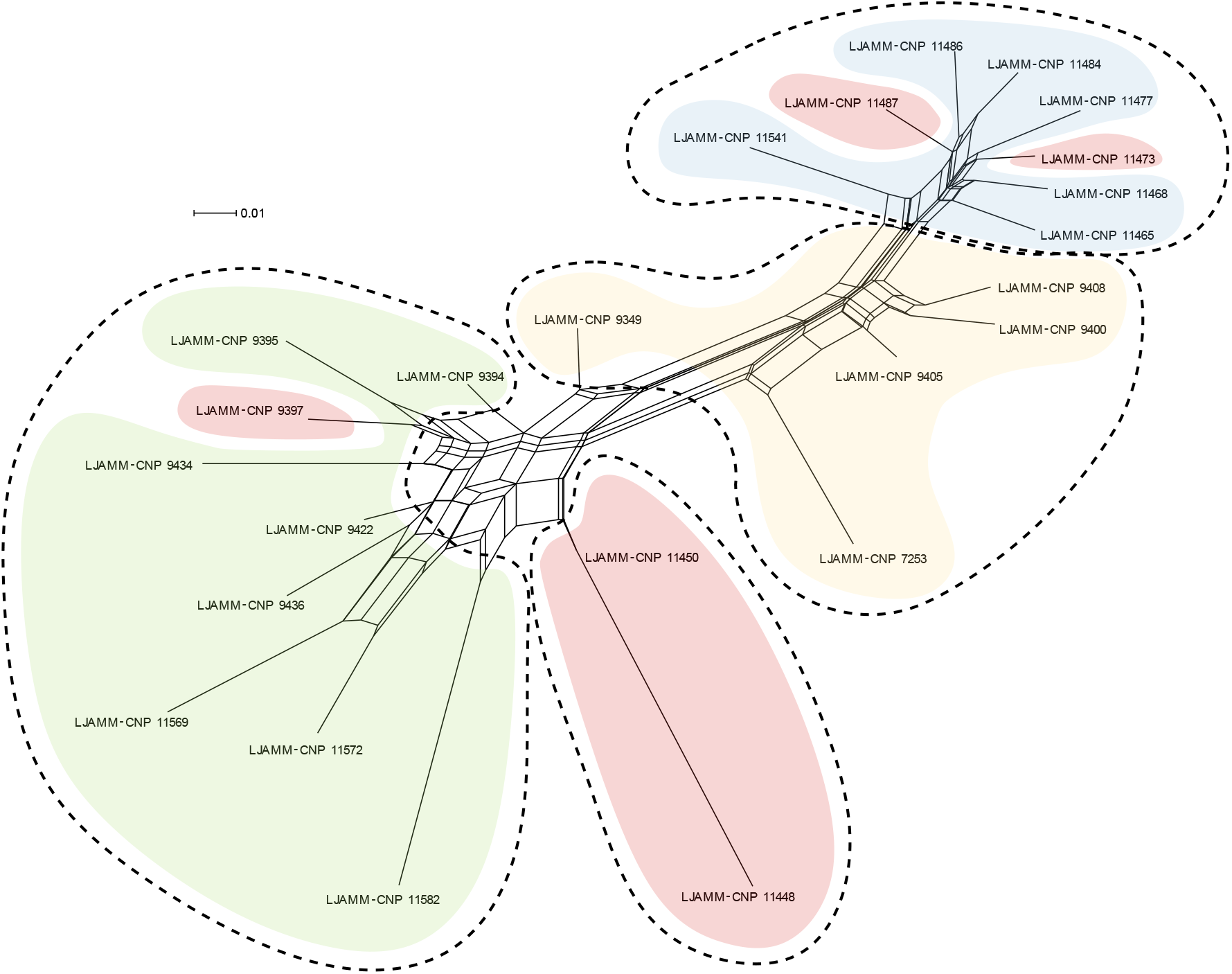
Phylogenetic network based on ddRADseq data using SplitsTree4. Clusters are colored according to the cyt-*b* haplogroups, and the dashed lines encircle the main clades recovered in the ddRADseq ML tree

## 4. Discussion

### 4.1. Lineage delimitation in the L. kingii group

It has been shown that some members of *Liolaemus* exhibit extensive levels of intraspecific variation, making diagnoses based on a single source of evidence unstable (e.g. coloration patterns, single-locus data; Breitman et al., 2013, 2015; Escudero et al., 2016; Esquerré et al., 2019). Further, a great number of species have been described based only on external morphological characters, not taking into account the extensive sexual and ontogenetic observed polymorphism, especially in the *L. kingii* group (see reviews in Breitman et al., 2013, and Avila et al., in press). Many instances of hybridization have also been detected in this genus (see Morando et al., 2020, for a recent review), which coupled with recent divergence times, could complicate species assignments, especially when a low number of molecular markers are employed (Medina et al., 2014). Taking these factors into account, we employed an integrative approach using genetic and morphological data to elucidate the relationships and species limits of the southernmost members of the *L. kingii* group.

The main phylogenetic relationships inferred with cyt-*b* and ddRADseq markers were concordant, and also with some previous phylogenetic hypotheses (Breitman et al., 2011, 2015; Pyron et al., 2013; Olave et al., 2014; Esquerré et al., 2018). The exception in our study was *L. tari*; this taxon was inferred as a well supported clade in the cyt-*b* tree, but paraphyletic in the ddRADseq tree (see Figs. 3, 6 and 7). Previous authors have postulated other alternative relationships, in particular for the positions of *L. baguali* and *L. sarmientoi* (Espinoza et al., 2004; Schulte and Moreno-Roark, 2010; Breitman et al., 2011; Olave et al., 2014). We also found incongruences between the cyt-*b* and ddRADseq trees with respect to the topological position of some individuals of *L. sarmientoi* (see section 4.2). Incongruence between different markers and phylogenetic reconstruction methods might reflect underlying evolutionary processes such as past or present hybridization and incomplete lineage sorting (Maddison, 1997; Knowles and Carstens, 2007; Degnan and Rosenberg, 2009). Further, these patterns are likely to be more easily detected in recently diverged populations or species, because a substantial amount of time is required, after the initial divergence of lineages, for multiple loci to sort to reciprocal monophyly with high probability (Knowles and Carstens, 2007).

Three of the four single-locus species delimitation approaches recovered the nominal species, with ABGD lumping *L. escarchadosi* + *L. tari* into a single entity. In contrast, *L*. sp. A was recovered as an independent lineage by all four methods (Fig. 3). The tree-based methods (i.e. mlGMYC, bGMYC and mPTP) tended to over-split the main haplogroups (6-13 entities), while the distance-based ABGD recognized fewer entities (4 entities). Previous studies have showed that tree-based species delimitation approaches tend to over-split entities (Dellicour and Flot, 2018). GMYC works well when the transition corresponding to diversification between species and genealogical branching within species is clearly demarcated (Reid and Carstens, 2012; Fujisawa and Barraclough, 2013; Blair and Bryson, 2017). However, this method is sensitive to sampling effort and general phylogenetic history, including divergence times, speciation rates, and differences in *Ne* among species (Reid and Carstens, 2012; Blair and Bryson, 2017). On the other hand, mPTP has proven to be consistent in delimiting putative species with varied sampling regimes and effective population sizes (Blair and Bryson, 2017). This suggests that this method may be suitable for a wide variety of empirical data sets because the error prone step of tree ultrametrization, which is necessary for the GMYC approach, is omitted by directly analyzing mutational steps along tree branches (Kapli et al., 2017; Eberle et al., 2019). Contrary to the tree-based approaches, the distance-based method ABGD is known to over-lump entities, especially when mutation rates are low and/or speciation rates are high (Dellicour and Flot, 2018). Taking into account the shortcomings inherent to these methods, taxonomic changes should not be made solely on these type of analyses; we follow recommendations to use these algorithms to generate primary species hypotheses (Leria et al., 2019), that can then be cross-compared with results based on other data sets (i.e. independent nuclear markers, morphometric data, etc.) following an integrative taxonomic workflow (Dayrat, 2005; Knowles and Carstens, 2007; Fujita et al., 2012; Talavera et al., 2013; Zhang et al., 2013; Sukumaran and Knowles, 2017; Dellicour and Flot, 2018; Fig. 1).

Our linear + meristic dataset resolved the highest number of differences between *L. baguali* and *L. escarchadosi*, two species that seem to be completely allopatric from each other (Fig. 2 and 4). Next, *L*. sp. A was partially distinguished, in concordance with the single-locus based delimitation, so it will be necessary to collect more individuals of this lineage and information of nuclear markers to test the boundaries of *L*. sp. A, and to compare it with the other members of the *L. kingii* group. Supporting the possible independent status of this lineage, Breitman et al. (2015) noted in-life coloration differences between *L*. sp. A and *L. tari*, *L. baguali*, *L. escarchadosi* and *L. sarmientoi*. *Liolaemus* sp. A is characterized by a pattern of small light dots scattered across the body with or without transverse bands (Breitman et al., 2015), whereas *L. tari* has a homogeneous brownish-gray color, and *L. baguali*, *L. escarchadosi* and *L. sarmientoi* share either dorsal transverse bands or a vertebral stripe. Linear, meristic and head shape data have been used to delimit some *Liolaemus* species (e.g. Minoli et al., 2016; Esquerré et al., 2019) and, in some cases, these characters proved to be good estimators. In this study, however, the partial level of resolution of these are relevant if one considers that *Liolaemus*, in general, has been characterized by morphological stasis, with many species sharing what appears to be a conserved morphology (Olave et al., 2017, 2018; Esquerré et al., 2019; Villamil et al., 2019). The evolution of morphological differences may reflect life history traits (e.g. habitat use, competition or sexual selection), thus, likely driven by more than one factor. Particularly for *Liolaemus*, the generalist morphology (Tulli et al., 2011) could provide advantages over the specialists in fluctuating and heterogeneous environments (Olave et al., 2020), especially in the complex landscapes of Southern South America (Rabassa et al., 2011).

*Liolaemus baguali* and *L. sarmientoi* were clearly differentiated with the population structure analyses, whereas the distinction between *L. escarchadosi* and *L. tari* was revealed only when they were analyzed separately, a result concordant with the inference that *L. tari* is paraphyletic with respect to *L. escarchadosi* in the ddRADseq tree (see Figs. 6 and 7). We note that these species are two of the most recently diverged within the *L. kingii* group (≈ 0.74 Mya; Esquerré et al., 2018). When a species originates, especially if it originates relatively quickly (as evidenced with the short branches that separate them), or on the periphery of another species’ range, the ancestral taxon will usually be rendered paraphyletic (Kuchta et al., 2018). However, over time, gene trees are expected to experience a transition to monophyly in a rate proportional to effective population sizes and rates of migration among populations (Baum and Shaw, 1995; Kuchta et al., 2018). Our study suggests that this and other *Liolaemus* species are at various points along this transition

The independent status of *L. escarchadosi* and *L. tari* was also corroborated by the coalescent-based BFD* (the top ranked model considers them as independent entities, see Table 3) and iBPP methods (PP = 1 for both lineages and with all datasets), and no clear signs of admixture was detected between them. Because the MSC assumes random mating (panmixia) within species (Barley et al., 2018), some limitations of the coalescent-based methods have been highlighted when intraspecific genetic structure is strong (Sukumaran and Knowles, 2017). Specifically, could result in the delimitation of “artefactual species” along boundaries that exhibit no barriers to gene flow (see Eberle et al., 2019, and Villamil et al., 2019 for discussions). This also raises the question that most current SDL methods based on this model do not account for introgressive hybridization (see below), isolation by distance (IBD) and ancient population structure (Barley et al., 2018). In the context of these “worst-cases scenarios” (Kuchta et al., 2018), such methods include simplifying assumptions that can lead them to fail with such complex patterns (Carstens et al., 2013; Kuchta et al., 2018). Nevertheless, the multispecies coalescent is the most objective approach available to explore species limits using genome-wide data (Fujita et al., 2012; Leaché et al., 2014, 2019; Villamil et al., 2019), and, given previous taxonomic treatments and geographical concordance, we are confident of the results obtained with these methodologies.

### 4.2. Introgression, incongruence between mitochondrial and ddRADseq markers and possible phylogeographic scenarios

Documented cases of hybridization between members of *Liolaemus* are now common, and could be a side-effect of the extraordinary diversification rates manifested by this genus (see detailed reviews in Olave et al., 2018, and Morando et al., 2020). Further, when coupled with widespread morphological stasis in some species occupying fluctuating environments, isolated populations may respond by: (i) rapidly increasing genetic variability after population bottlenecks, and/or (ii) contributing to the maintenance of an adaptive generalist biology by rapidly generating a wide distribution of intermediate phenotypes in cohorts of hybrid descendants (Olave et al., 2018).

Particularly for the *L. kingii* group, Breitman et al. (2011) hypothesized an ancient hybridization event that produced a lineage of uncertain taxonomic status (*L*. sp. 4), given its ambiguous phylogenetic placement with respect to other members of the group (based on mitochondrial and nuclear data); this result was confirmed by Olave et al. (2014). Most important, morphological signatures of hybridization were also detected for the species studied in this work (Breitman et al., 2015), specifically for: *L. baguali × L. sarmientoi, L. baguali* × *L. tari*, and *L. escarchadosi* × *L. tari*. Here, the population structure analyses corroborate the previous findings. fastSTRUCTURE plots revealed signs of admixed genomes in specimens LJAMM-CNP 11450 (assigned to *L. sarmientoi*), LJAMM-CNP 9349 and LJAMM-CNP 7253 (both assigned to *L. tari*), suggesting recent hybridization for *L. baguali* × *L. sarmientoi*, and *L. baguali* × *L. tari* (see Fig. 6a, b). In addition, *D*-statistic tests revealed weak signals of introgression between these lineages, and interestingly for *L. sarmientoi* × *L. tari* (Table 4), although these last two species revealed no signs of admixture with fastSTRUCTURE nor DAPC. In this latter case, it would be necessary to collect more data from both species to distinguish between real hybridisation/introgression vs. a sample size artifact. The individual LJAMM-CNP 11450 was collected in the periphery of the distribution of *L. sarmientoi* (locality 8), far from the distribution of *L. baguali*, but since the *D*-statistic test detected a weak introgression signal between them, we could argue that ancient hybridization events could have been responsible. In the same line of reasoning, one of the admixed individuals of *L. tari* (LJAMM-CNP 9349) was sympatric with members of *L. baguali* (in locality 40), suggesting ongoing or recent hybridization between these species. We are aware of our limited genomic sampling, but clarification will require more sampling over a larger geographic region. Future studies also need to explore whether a generalist morphology and apparent phenotypic stasis in *Liolaemus* might facilitate the ongoing gene flow among divergent lineages.

An interesting observation that three individuals identified as *L. sarmientoi* in the cyt-*b* gene tree were grouped within *L. baguali* (specimen LJAMM-CNP 9397) and *L. escarchadosi* (specimens LJAMM-CNP 11473 and LJAMM-CNP 11487) in the ddRADseq tree (see Figs. 6, 7 and 9). The individual LJAMM-CNP 9397 was sympatric with *L. baguali* at locality 38, and previous observations by Breitman et al. (2015) noted that this specimen has a dorsal stripe pattern more similar to *L. baguali* than to *L. sarmientoi*. On the other hand, the specimens LJAMM-CNP 11473 and LJAMM-CNP 11487 also identified as *L. sarmientoi* were sympatric with *L. escarchadosi* in localities 19 and 22, with the caveat that, given the extensive morphological variation within each species, could not been morphologically assigned to either *L. sarmientoi* or to *L. escarchadosi* by Breitman et al. (2015). We do not consider incomplete lineage sorting as most plausible because none of the species correspond to sister lineages (*L. sarmientoi* with *L. baguali*, and *L. sarmientoi* with *L. escarchadosi*), otherwise we could expect a high probability of incompatibilities in gene trees.

The high levels of introgression detected here could be partly explained by the high level of sympatry between most of the lineages in several localities (Fig. 2). *Liolaemus baguali* has a north-western distribution along the Asador Plateau and adjacent areas (north to Viedma Lake), and in some localities it is sympatric with *L. sarmientoi* (localities 38, 39 and 41) and *L. tari* (locality 40). *Liolaemus tari*, at the same time, is sympatric with *L. escarchadosi* in its type locality (37) and *L*. sp. A (locality 34), which in turn, is sympatric with *L. escarchadosi* (locality 31). *Liolaemus tari* and *L*. sp. A have more restricted distributions on the western side of Santa Cruz province. *Liolaemus sarmientoi* has an eastern distribution, peripheral to *L. escarchadosi*, but the two are sympatric in the north (locality 6), east (localities 14 and 15), and south (localities 19 and 22). The species that seem to be completely allopatric to each other are *L. baguali* with respect to *L. escarchadosi*, and *L. sarmientoi* with respect to *L*. sp. A and *L. tari*. Contact areas offer opportunities for past or present hybridization and introgression, especially in closely related or recently diverged species, where reproductive isolation might not be complete (Mallet, 2007), or among non-sister species as well during rapid adaptive radiations (Mallet et al., 2016). It is interesting to note that contact areas between the above species are mostly distributed in the western region, along the eastern flank of the Andes (between 49°S and 51°S), and mainly in the areas surrounding the Viedma and Argentino Lakes (Fig. 2). This area has been identified as a potential phylogeographic refugium (i.e. *in situ* persistence during Pleistocene glaciations) for numerous plant genera, and populations of the *L. lineomaculatus* group (Sérsic et al., 2011; Breitman et al., 2012). During glacial advances, much of the original extra-Andean landscape cooled to form permafrost, so the *in situ* formation of periglacial refugia exposed suitable habitats for organisms to shift their distributions to less harsh climates (Breitman et al., 2012). This series of events could have promoted repeated rounds of secondary contact between diverging lineages. In addition, the geological evidence present in this area suggested the occurrence of at least nine glacial advances from the Pliocene to the Last Glacial Maximum (LGM = 0.025–0.016 Mya) (Rabassa et al., 2011). It is well established that Pleistocene glacial cycles promoted the contraction and subsequent expansion of lineage distributions (Hewitt, 1996), which in turn forced the reproductive isolation during interglacial periods but fostered secondary contact during glacial periods (e.g. Fuentes and Jaksić, 1979). It is also interesting to note the patterns of genetic structure relative to the presence of natural barriers, such as the break in the Chalía-Chico river systems (Fig. 2). This was evident mainly in the result from DAPC analysis (Fig. 6d), which shows broad northern (*L. baguali* + *L. sarmientoi*) and southern clusters (*L. escarchadosi* + *L. tari*). The Chico River basin has been regarded as a phylogeographic break in previous studies on herbs (*Calceolaria polyrhiza*; Cosacov et al., 2010) and lizards of the *L. lineomaculatus* group (Breitman et al., 2012). Fragmented habitats influenced relatively recent patterns of diversification in several taxa in southern Patagonia (see Sérsic et al., 2011, and Breitman et al., 2012 for a summary), and this work constitute another example for the observed patterns.

## 5. Concluding remarks

In this work we took advantage of recently developed statistical approaches and the availability of different datasets to infer species limits in the southernmost members of the *L. kingii* group. We highlight three important outcomes of this study. First, we confirmed the validity of the species *L. baguali, L. tari, L. escarchadosi* and *L. sarmientoi* with an integrative approach, and refined their geographic distributions. Second, we provided with more evidence for the possible independent status of *L*. sp. A; we empirically demonstrated the value of simultaneous analyzes of different types of data to test explicit species hypotheses, with the potential to overcome problems created by relying on a either a single source of evidence or a single method, to infer species limits. Finally, we found additional evidence of introgressive hybridization between species of *Liolaemus*, specifically for *L. baguali* × *L. sarmientoi*, *L. baguali* × *L. tari*, and *L. sarmientoi* × *L. tari*; corroborating and extending early suggestions of Breitman et al. (2015).

## Data accessibility

New cytochrome-*b* data are available at NCBI’s GenBank with accession numbers xxx-xxx. ddRAD data (raw individual sequence reads) are available at NCBI’s Short Read Archive (SRA) under bioproject xxx.

## Conflict of interest

We, the authors, declare no competing interests.

## Acknowledgments

We are grateful to past and present members of the Grupo de Herpetología Patagónico (GHP) for assistance in field collections, animal curation and laboratory procedures. We thank Adam D. Leaché for facilitating ddRADseq data collection. We also thank E.G. Díaz-Huesa for helpful comments on the manuscript and J. Villamil and D. Esquerré for advice with the ddRADseq data assembly. Financial support was provided by a CONICET graduate fellowship (K.I.S.) and the following grants: FONCYT PICT-2011-0784 (issued to L.J.A.), CONICET-PIP 0336/13, FONCYT-PICT-2011-1397 and ANPCYT-FONCYT 1252/2015 (issued to M.M.), and NSF-PIRE (OISE 0530267) and Macrosystems (EF 1241885) awards. We thank the fauna authorities from Santa Cruz Province for collection permits.

## AppendixA. Supplementary material

Supplementary data to this article can be found online at

## Notes

### Competing Interest Statement

The authors have declared no competing interest.

